# The Y chromosome contributes to sex-specific aging in Drosophila

**DOI:** 10.1101/156042

**Authors:** Emily J. Brown, Alison H. Nguyen, Doris Bachtrog

## Abstract

Heterochromatin suppresses repetitive DNA, and a loss of heterochromatin has been observed in aged cells of several species, including humans and Drosophila. Males often contain substantially more heterochromatic DNA than females, due to the presence of a large, repeat-rich Y chromosome, and male flies generally have shorter average life spans than females. Here we show that repetitive DNA becomes de-repressed more rapidly in old male flies relative to females, and repeats on the Y chromosome are disproportionally mis-expressed during aging. This is associated with a loss of heterochromatin at repetitive elements during aging in male flies, and a general loss of repressive chromatin in aged males away from pericentromeric regions and the Y. By generating flies with different sex chromosome karyotypes (XXY females; X0 and XYY males), we show that repeat de-repression and average lifespan is directly correlated with the number of Y chromosomes. Thus, sex-specific chromatin differences contribute to sex-specific aging in flies.

## Introduction

The chronic deterioration of chromatin structure has been implicated as one of the molecular signatures of aging (O’Sullivan and Karlseder 2012; Tsurumi and Li 2012; Wood, et al. 2010), and an overall loss of heterochromatin and repressive histone marks is observed in many old animals (Haithcock, et al. 2005; Larson, et al. 2012; Zhang, et al. 2015). Heterochromatin is enriched at repetitive DNA, and its loss can result in de-repression and mobilization of silenced transposable elements (TEs) (De Cecco, et al. 2013a; De Cecco, et al. 2013b; Elsner, et al. 2018; Li, et al. 2013; Van Meter, et al. 2014; Wood and Helfand 2013; Wood, et al. 2016). The amount of repetitive DNA can differ substantially between sexes, due to the presence of a highly repetitive (and normally poorly assembled) Y or W chromosome in the heterogametic sex. In the fruit fly *Drosophila melanogaster*, for example, males contain a ~40-Mb large completely repetitive Y chromosome (Chang and Larracuente 2019; Hoskins, et al. 2002), while the pericentromeric heterochromatin on the X only amounts to ~10-20-Mb (depending on the strain). Thus, this implies that a substantially larger fraction of the male genome is heterochromatic compared to the female genome (Hoskins, et al. 2002). Males have a shorter average lifespan in many taxa, including humans and most Drosophila species (Lehtovaara, et al. 2013; Tower and Arbeitman 2009; Yoon, et al. 1990). Indeed, the genetic sex determination system predicts adult sex ratios in tetrapods, with the heterogametic sex being less frequent (Pipoly, et al. 2015). Lower survivorship of the sex with the repetitive Y or W chromosome may suggest a link between sex-specific mortality, chromatin and sex chromosomes.

Here, we test for an association between sex-specific heterochromatin loss and a de-repression of repetitive DNA during aging, by assaying chromatin and gene expression profiles in young and aged individuals of *D. melanogaster*. We further create flies with different sex chromosome karyotypes (that is, X0 and XYY males and XXY females), to directly test for the influence of the Y chromosome on longevity and sex-specific changes in gene expression.

## Results

### Drosophila males and females differ in repeat content and longevity

We chose *D. melanogaster* to investigate the contribution of the Y chromosome to sex-specific aging, since a substantial fraction of its heterochromatin, including its Y chromosome, has been assembled (Hoskins, et al. 2015). In addition, the availability of mutant strains allows us to generate flies that differ in their sex chromosome configuration. **Figure 1A** shows a schematic overview of the karyotype for *D. melanogaster* males and females, and the approximate size and position of large heterochromatic segments (Hoskins, et al. 2002). Previous studies have suggested that males have approximately 20Mb more heterochromatin per cell than females (**Figure 1A**) (Hoskins, et al. 2002), and flow cytometry estimates confirm that the Canton-S strain used here shows similar differences in repeat content between the sexes (**Table 1**). Thus, this confirms that male Canton-S flies contain significantly more repetitive DNA than female.

**Figure 1.**
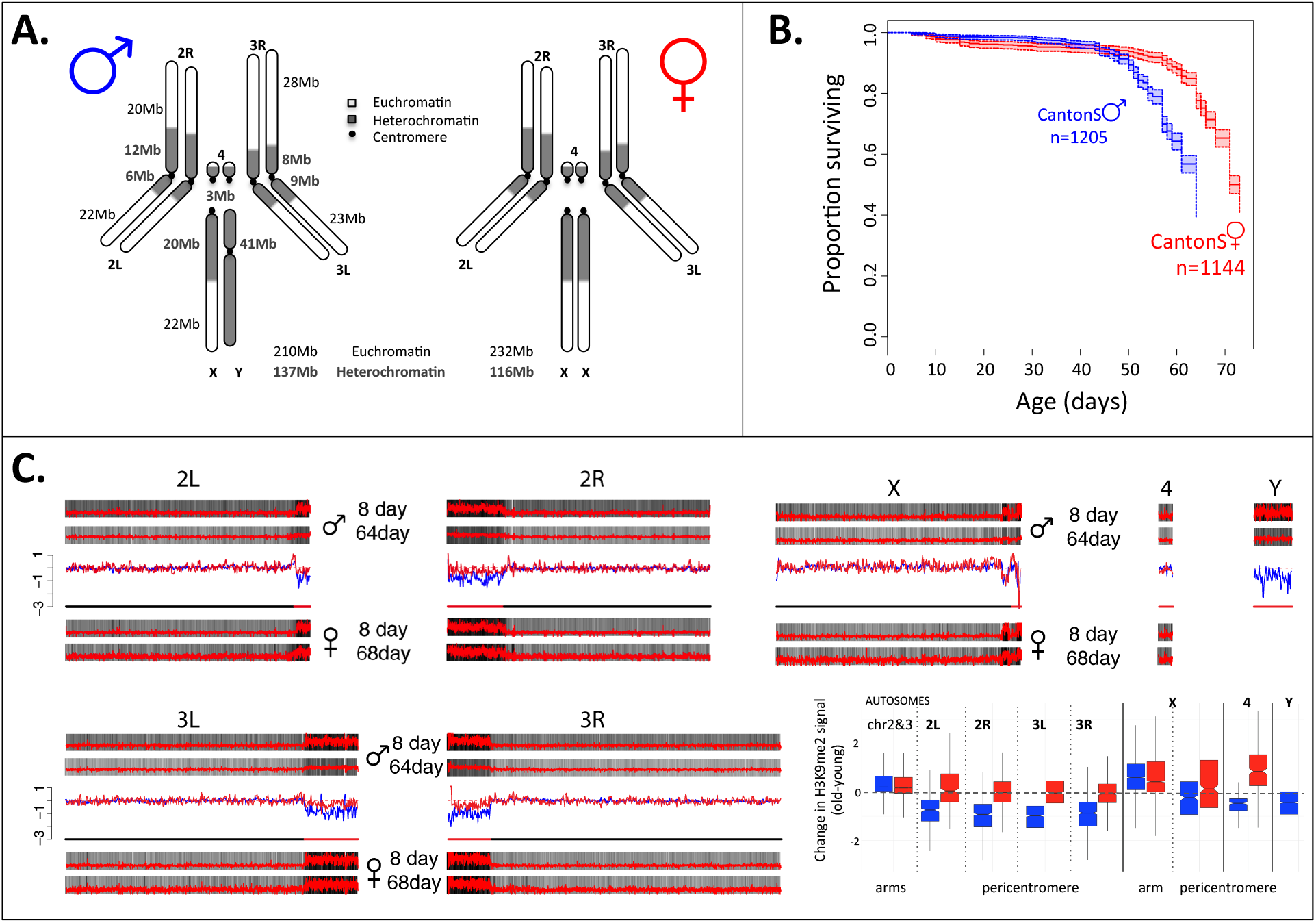
Aging and the sex-specific chromatin landscape in Drosophila. **A.** Kaplan-Meier survivorship curves (Kaplan and Meier 1958) for Canton-S males (blue) and females (red), with the shaded region indicating the upper and lower 95% confidence interval calculated from the Kaplan-Meier curves. Karyotypes of male and female *D. melanogaster* are shown, with heterochromatic regions indicated in blue, and euchromatic regions in gray. The number of flies counted for each sex (n) to obtain the survivorship curves is indicated. **B.** Genome-wide enrichment of H3K9me2 for young (8 days) and old (64 or 68 days) *D. melanogaster* males and females along the different chromosome arms. Enrichment in 5kb windows is shown in red lines (normalized ratio of ChIP to input, see Materials & Methods), and the enrichment in 20kb windows is shown in gray scale according to the scale in the lower right of the panel, with the darkest gray corresponding to the highest 5% of values across all windows from all samples, and the lightest gray corresponding to the lowest 10% of values across all windows from all samples. Subtraction plots show the absolute difference in signal of 50kb windows between young and aged flies along the chromosome arms, with each sample further smoothed by subtracting out the median autosomal euchromatin signal, with females in red and males in blue, and the pericentromeric region of each chromosome indicated by the red segment of the line beneath each chromosome. **C.** Box plot showing the smoothed ChIP signal for all 5kb windows in different chromosomal regions (* p<0.05, ** p<1e-6, *** p<1e-12, Wilcoxon test) for males (blue) and females (red), with pericentromere boundaries defined by the Release 6 version of the *D. melanogaster* genome.

**Table 1.**
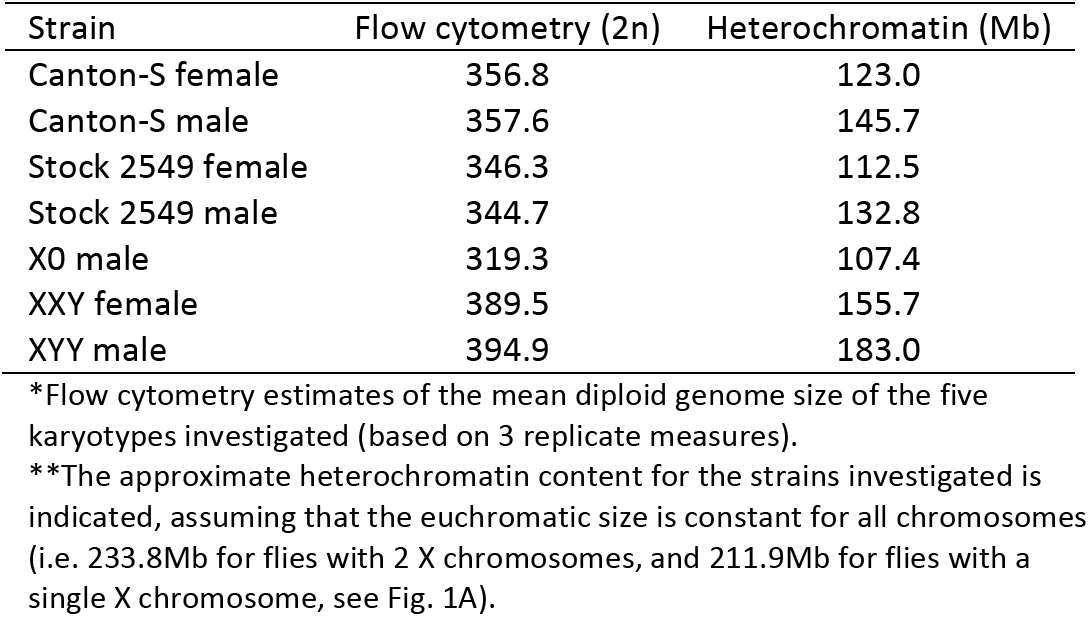
Flow cytometry estimates of the mean diploid genome size

Previous research in Drosophila showed that for the vast majority of species, male flies live significantly shorter than female flies (Yoon, et al. 1990). In particular, a study surveying longevity in 68 species (and 89 strains) of Drosophila found statistically significant differences in longevity between the sexes for 55 strains. Females had higher longevity in 53 strains, and males lived longer in only 2 strains (Yoon, et al. 1990). We determined longevity for males and females of two standard lab strains from *D. melanogaster* (Canton-S and Oregon-R), and the 2549 strain from the Bloomington Stock Center, which has a compound metacentric X chromosome (that is, two X chromosomes fused at the centromere) and a heterocompound X-Y chromosome (that is, an X chromosome inserted between the two arms of the Y chromosome). Lifespan assays confirm that males live significantly shorter than females, for both wildtype strains and the 2549 strain (**Figure 1B**, see **Figure S1** for lifespan assays in Oregon-R and 2549). Increased longevity of females is consistent with multiple studies on sex-specific lifespan in Drosophila (Tower and Arbeitman 2009; Yoon, et al. 1990).

### Heterochromatin loss differs between sexes

We gathered replicate ChIP-seq data for a repressive histone modification typical of heterochromatin (H3K9me2) from young 8-day and old 64-68-day *D. melanogaster* males and females (Canton-S), to test for sex-specific heterochromatin loss during aging. We used a ‘spike in’ normalization method to compare the genomic distribution of chromatin marks across samples (Brown, et al. 2019; Li, et al. 2014). Specifically, we spiked-in a fixed amount of chromatin from *D. miranda* to each *D. melanogaster* chromatin sample prior to ChIP and sequencing. We chose this species to serve as an internal standard for our *D. melanogaster* ChIPs, since they are sufficiently diverged from each other that there is very little ambiguity in the assignment of reads to the correct species. We employed a previously described normalization strategy (Li, et al. 2014), where the relative recovery of *D. melanogaster* ChIP signal vs. *D. miranda* ChIP signal, normalized by their respective input counts, was used to quantity the relative abundance of the chromatin mark in *D. melanogaster*. Note that this normalization strategy accounts for differences in ploidy levels of sex chromosomes (see **Figure S1** in ref. Brown, et al. 2019). We also used a linear regression model to estimate the relative recovery of *D. melanogaster* ChIP signal vs. *D. miranda* ChIP signal, normalized by their respective input counts (Bonhoure, et al. 2014). Overall enrichment patterns and differences between sexes and ages are quantitatively similar between the two methods, showing that our inferences are robust to our normalization strategy (**Figure S2A**).

Repetitive regions pose a challenge for mapping with short reads, since one cannot be sure that a particular locus is generating the reads in question if they map to multiple positions. Our study is concerned with the overall behavior of repetitive regions in the genome during aging, and not focused on any particular locus. Thus, analyzing all reads (including those mapping to multiple locations) is most appropriate for our purpose. However, we repeated our analysis using only uniquely mapping reads, which confirms that our inferences are robust when only considering uniquely mapping reads (**Figure S2B**).

**Figure 1C** shows the genomic distribution of the repressive histone modification H3K9me2 for young and old male and female flies. As expected, heterochromatin is enriched at repetitive regions, including pericentromeres, the small dot chromosome and the repeat-rich Y. While the genomic distribution of H3K9me2 looks similar between young males and females (Brown, et al. 2019), heterochromatin enrichment changes dramatically in old male but less so in old female flies. In particular, we see a general loss of heterochromatin at repetitive regions in aged males (**Figure 1C; Figure S3** for biological replicate), and males show significantly more regions that lose H3K9me2 signal (1.5-fold or more) during aging compared to females (232 vs. 73, p<2.2e-16, Fisher’s exact test; **Figure S4**). Almost all regions that lose heterochromatin are located within the pericentromere or the Y chromosome (**Figure S4,S5**). On the other hand, fewer regions in males gain H3K9me2 signal (1.5-fold or more) during aging relative to females (6 vs. 120, p<2.2e-16, Fisher’s exact test; **Figure S4**). Genomic regions that gain H3K9me2 signal are enriched on the X of females (p<2.2e-16, Fisher’s exact test; **Figure S4,S5**), and tend to be located close to the pericentromeric boundary (p=0.04 Fisher’s exact test; **Figure S6, Figure S4,S5**). This suggests that heterochromatin / euchromatin boundaries are less efficiently maintained in old flies, resulting in spreading of heterochromatin from the repeat-rich pericentromere into neighboring regions.

Thus, our results show that male flies lose heterochromatin marks more rapidly than female flies in *D. melanogaster* and our results are reproducible using different mapping and normalization strategies, and independent biological replicates.

### Mis-expression of heterochromatic genes in old flies

Sex-specific chromatin changes during aging are associated with sex-specific expression changes. To study gene expression during aging, we gathered replicated stranded RNA-seq data from young and old flies. We find that genes located in pericentromeric regions change their expression more during aging compared with genes in chromosomal arms, in both sexes (**Figure S7**). While heterochromatin typically has a repressive effect on gene expression, genes located in normally heterochromatic regions (such as the pericentromere) are known to depend on this repressive chromatin environment for proper transcription (Lu, et al. 2000). Indeed, the global loss of heterochromatin in pericentromeric regions is associated with reduced expression levels of pericentromeric genes in aged males and females (**Figure S7**). Genes that gain the H3K9me2 mark during aging tend to decrease in expression (**Figure S8,S9**).

Overall, we find that gene expression during aging differs between males and females. Of the top 10% of genes that are differentially expressed during aging in males and females, 35.5% show expression changes in both sexes (**Figure S10**). Chromosomal location does not appear to be the main determinant influencing gene expression during aging; of the 464 genes that are most differentially expressed in both males and females, only 17 are located inside or within 1Mb of the pericentromere (we expect 22 by chance). The most differentially expressed genes affect similar functional categories in males and females (including GO categories “antibacterial humoral response”, “macromolecule biosynthetic process”, “reproduction”, and “translation”, **Figure S11**), but many GO terms are unique to one sex (13 highly significantly enriched (p<10-9) GO terms shared by both sexes; 13 male-specific and 1 female-specific GO term). The rDNA locus has also been shown to be mis-expressed during aging (Larson, et al. 2012); we generated rDNA-depleted libraries, which precludes us from studying rDNA expression.

### Repeat de-repression in old male flies

In addition, we find sex-specific differences in repeat reactivation during aging. We mapped our transcriptome data to the consensus repeat library of *D. melanogaster* and detect low levels of expression of repetitive elements in young male and female flies (**Figure 2A**). Aged females maintain efficient repression of TEs, while expression for the major classes of annotated TEs increases during aging for males (**Figure 2A,C**). De-repression of TEs is more pronounced in males both in terms of the number of individual elements that show a significant increase in expression during aging, as well as the fraction of the transcriptome that consists of repetitive transcripts across all repetitive elements.

**Figure 2.**
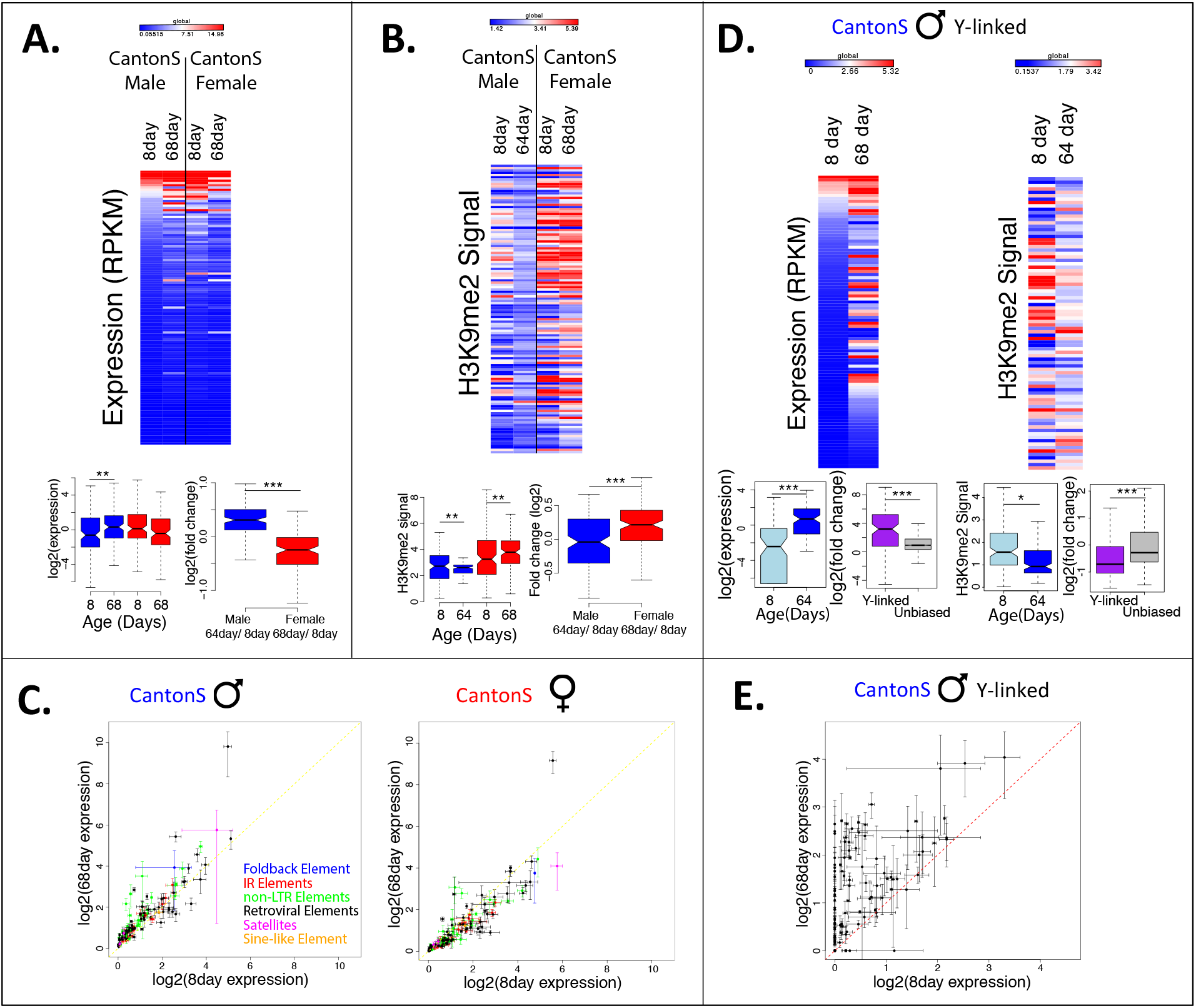
Sex-specific silencing and expression of repeats during aging. **A.** Expression of all repeats from FlγBase consensus library from Release 6 of the *D. melanogaster* genome in young (8day) and old (~68day) male and female Canton-S, averaged across replicates, with significance values calculated using the Wilcoxon test (* p<0.05, ** p<0.01, *** p<1e-5). Heatmaps are visualized globally according to the scale, with dark red corresponding to the top 5% of all values across all samples and dark blue corresponding to the bottom 5% of all values across all samples. **B.** H3K9me2 enrichment in repeats from FlyBase consensus library in young (8day) and old (64 or68day) male and female Canton-S, averaged across replicates, with significance values calculated using the Wilcoxon test (* p<0.05, ** p<0.01, *** p<1e-5). The heatmap is scaled in the same manner as in (A.). **C.** Expression for repeat families for old and young males and females, with lines indicating the standard deviation for each estimate of expression across replicates and colors indicating the class of repetitive element. **D.** Expression and H3K9me2 signal in putatively Y-linked repeats in young (8day) and old (64 or 68day) Canton-S males, with significance values calculated using the Wilcoxon test (* p<0.01, ** p<1e-4, *** p<1e-10). Heatmaps are scaled in the same manner as in (A). **E.** Expression of putatively Y-linked repeats for old and young Canton-S males, with lines indicating the standard deviation for each estimate of expression across replicates.

Overall, we find that in females, 6 repetitive elements show a significant increase in expression during aging and 14 a significant decrease (**Figure 2C**), but the total fraction of transcripts derived from repeats increases during aging (the fraction of repetitive reads in all RNA-seq reads is 2.0% at 8 days, vs. 4.6% at 68 days, **Table S1, Figure S12**). The increase in repeat expression is much more pronounced in males, with 32 repetitive elements showing a significant increase in expression during aging and 4 showing a significant decrease (**Figure 2C, Figure S12**), and the total fraction of repetitive reads increases from 1.6% to 5.8% (**Table S1**). The TE showing the highest level of de-repression in both sexes is *copia*, which is expressed 28-fold more in old versus young males, and expressed 15-fold more in old versus young females (**Figure 2C**). H3K9me2 profiles at TE families show that there is a general enrichment of this repressive mark in young male and female flies (**Figure 2B**). Consistent with genome-wide expression profiles showing overall efficient silencing of repeats in old females, there is no global loss of the repressive chromatin mark at repetitive elements in 68-day old females (in fact, there is a slight increase, **Figure 2B**). However, aged *D. melanogaster* males undergo a general loss of the H3K9me2 histone modification in repetitive elements (**Figure 2B**). Thus, chromatin and gene expression profiles show that TEs lose their epigenetic silencing and become mis-expressed in old male flies.

Males have approximately 20% more repetitive sequence than females, due to the repeat-rich Y chromosome. The sex-specific increase in repeat expression may be triggered by the presence of the heterochromatic Y chromosome in males, and the Y indeed shows a dramatic loss of heterochromatin during aging (**Figure 1**). To see if Y-linked repeats are especially prone to mis-regulation during aging, we used *de novo* assembled male-specific and male-biased (putatively Y-linked) repetitive sequences (**Figure S13**) (Brown, et al. 2019). Indeed, we find that in males, putatively Y-linked repeats are up-regulated more strongly during aging, relative to repeats present in both sexes (p=5.9e-12, Wilcoxon test, **Figure 2D,E**). Overall, we find that 42 Y-linked repeats show a significant increase in expression during aging (and only one a significant decrease; **Figure 2D,E, Figure S14**), and the total fraction of transcripts derived from Y-linked repeats increases more than 9-fold in old males (**Table S2**).

Additionally, putatively Y-linked repeats disproportionately lose the repressive histone modification H3K9me2 during aging compared to other repeats (p=3.4e-11, Wilcoxon test, Figure 2D). Thus, male-biased and male-specific repeats, i.e. repeats that are located on the Y chromosome, are especially prone to de-repression during aging in males.

### Additional Y chromosomes decrease lifespan

To directly test whether the Y chromosome contributes to sex-specific TE de-repression and aging, we generated *D. melanogaster* females containing a Y chromosome (XXY flies), and males with either zero or two Y chromosomes (X0 and XYY flies), by crossing Canton-S flies to different strains with attached-X and attached-X-Y chromosomes (**Figure 3A**, see **Methods** for strain information). Note that X0 males and XXY females of a given cross have the same autosomal background, but differ in their genomic background from other crosses and from Canton-S, which can contribute to lifespan variation among strains (**Figure 3B,C**). Utilizing different strains to generate flies with aberrant sex chromosomes, however, should control for genomic background effects. Indeed, we find qualitatively identical results using three independent strains to generate XO/XXY/XYY flies, suggesting that differences in longevity are not due to genomic background, but caused by the presence or absence of the Y.

**Figure 3.**
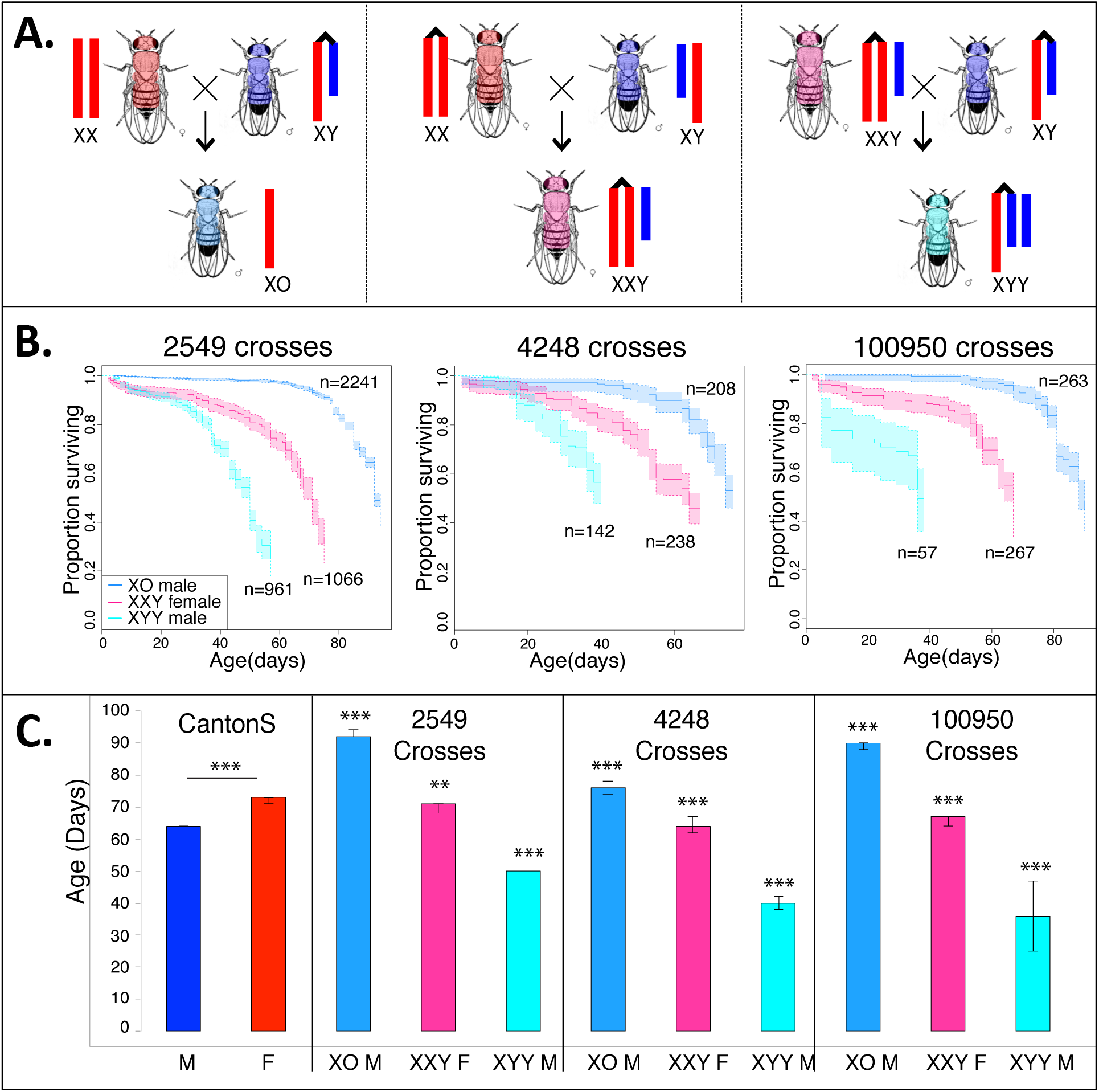
Survivorship of XXY females and X0 and XXY males. **A.** Schematic crossing scheme used to generate flies with aberrant sex chromosomes, with Canton-S used as the wild-type (wt) males and females for all crosses, and various lines with C(1)RM and C(1;Y) indicated by the attached X/ XY karyotypes. Crosses between wt females and attached XY males result in XO flies (left), crosses between attached X females and wt males result in XXY females (middle), and crosses between XXY females and attached XY males result in XYY males (right). **B.** Kaplan-Meier survivorship curves for flies with aberrant sex chromosome karyotype, generated with various C(1)RM and C(1;Y) lines as indicated at the top of each survivorship curve (stock 2549 and 4248 were obtained from the Bloomington Stock Center, and stock 100950 was obtained from Kyoto). The crosses to obtain the XO, XXY and XYY flies are shown in panel A. Shaded areas indicate the upper and lower 95% confidence interval calculated from the Kaplan-Meier curves. **C.** Median lifespan for each of the different karyotypes measured, with error bars indicating the upper and lower 95% confidence intervals (estimated by the Kaplan-Meier curves). Significance is compared to the wild-type Canton-S of the same sex for each aberrant karyotype, and was calculated using the survdiff package in R (* p<0.01, ** p<1e-6, *** p<1e-12).

In particular, we compared sex-specific lifespans of wildtype Canton-S *D. melanogaster* flies, and XXY female and XO and XYY male resulting from crosses to the three different attached-X and attached-X-Y strains (**Figure 3B,C**). Cumulative survival probabilities show that life span of females that contain a Y chromosome (XXY females) is reduced relative to wildtype females or males that lack a Y chromosome (X0 males) for all crosses assayed (**Figure 3B,C**). Indeed, X0 males show a dramatic increase in life span relative to wildtype males, and even outlive wildtype females (**Figure 3B,C**). X0 males are sterile and have the least amount of repetitive DNA of all karyotypes investigated (~10Mb less than Canton-S females and ~40Mb less than Canton-S males; **Table 1**); both of these factors may contribute to increased lifespan. Males with two Y chromosomes (XYY), in contrast, live the shortest (**Figure 3B,C**), and their lifespan is reduced considerably relative to wildtype males, despite both karyotypes being fertile. Thus, survivorship data suggest that the number of Y chromosomes directly influences organismal survival in Drosophila.

### Mis-expression of Y genes and repeats in flies with additional Y chromosomes

Gene expression changes during aging in the aberrant karyotypes show many of the same patterns as wildtype flies, with similar networks of GO terms enriched in both XO and XYY males, and XXY females, including “reproduction” or “sensory perception of chemical stimulus” (**Figure S11**). Overall, we find that 101 of the top 10% of genes mis-expressed during aging are shared among all 5 karyotypes (p<<1e-5, permutation test). These genes do not show any enrichment for a particular GO term, and only 6 of them are located inside or within 1Mb of the pericentromere (expect 4.8 genes).

Genomic location also influences gene expression changes during aging in flies with aberrant karyotypes. As in wildtype males and females, genes located in the pericentromere show a decrease in expression during aging in XXY females and X0 males (**Figure S15**; XYY flies show no significant expression change at pericentromeric genes). Y-linked genes in wild-type males are expressed almost exclusively in male reproductive tissues (Carvalho, et al. 2009), and we do not detect any expression of Y-linked genes even in very old XY male heads (**Figure S15**). In contrast, we find that Y-linked genes are inefficiently silenced in heads of XXY females and XYY males and are becoming de-repressed even more during aging (6 Y genes are expressed in old XXY females, and 14 in old XYY males, **Figure S15**).

Thus, wildtype males maintain efficient silencing at their Y-linked genes during aging, despite global heterochromatin loss on the Y chromosome and mis-expression of Y-linked repeats. Silencing mechanisms on the Y chromosome of XXY and XYY flies, on the other hand, appear to be generally compromised, even in young individuals. Indeed, we previously showed that the Y chromosome affects global heterochromatin integrity (Brown, et al. 2019). Young flies with additional Y chromosomes (XXY females or XYY males) show lower levels of H3K9me2 enrichment at their TEs, and a de-repression of Y-linked repeats relative to wildtype flies, while X0 flies showed increased levels of H3K9me2 at repeats (Brown, et al. 2019). Expression profiles in XXY females and XYY/X0 males demonstrate that the absence or presence of the Y chromosome modulates expression of TEs during aging (**Figure 4**). Expression profiles from aged flies with aberrant sex chromosome karyotypes confirm our expectation that X0 males show less de-repression of TEs during aging relative to wildtype males (**Figure 4A**). XXY females, on the other hand, show more mis-expression of repeats during aging compared to wildtype females. In XXY females, 7 elements show a significant increase in expression during aging and 3 elements show a significant decrease in expression (**Figure 4A**, compared to 6 /14 elements that increase/decrease expression in wild-type females), and the fraction of repetitive transcripts increases more during aging for XXY females (3.1-fold increase in XXY females vs. 2.2-fold in wildtype females, **Table S1**). XYY males show the greatest number of repeats with significantly increased expression during aging (33 elements, **Figure 4A**), but not the highest fold change in total fraction of repetitive reads (**Table S1**), partly because young XYY males already show the highest expression of repeats of any of the 5 karyotypes (**Table S1**), and partly because old XYY males are approximately 30 days younger than the other karyotypes (**Figure 3B**).

**Figure 4.**
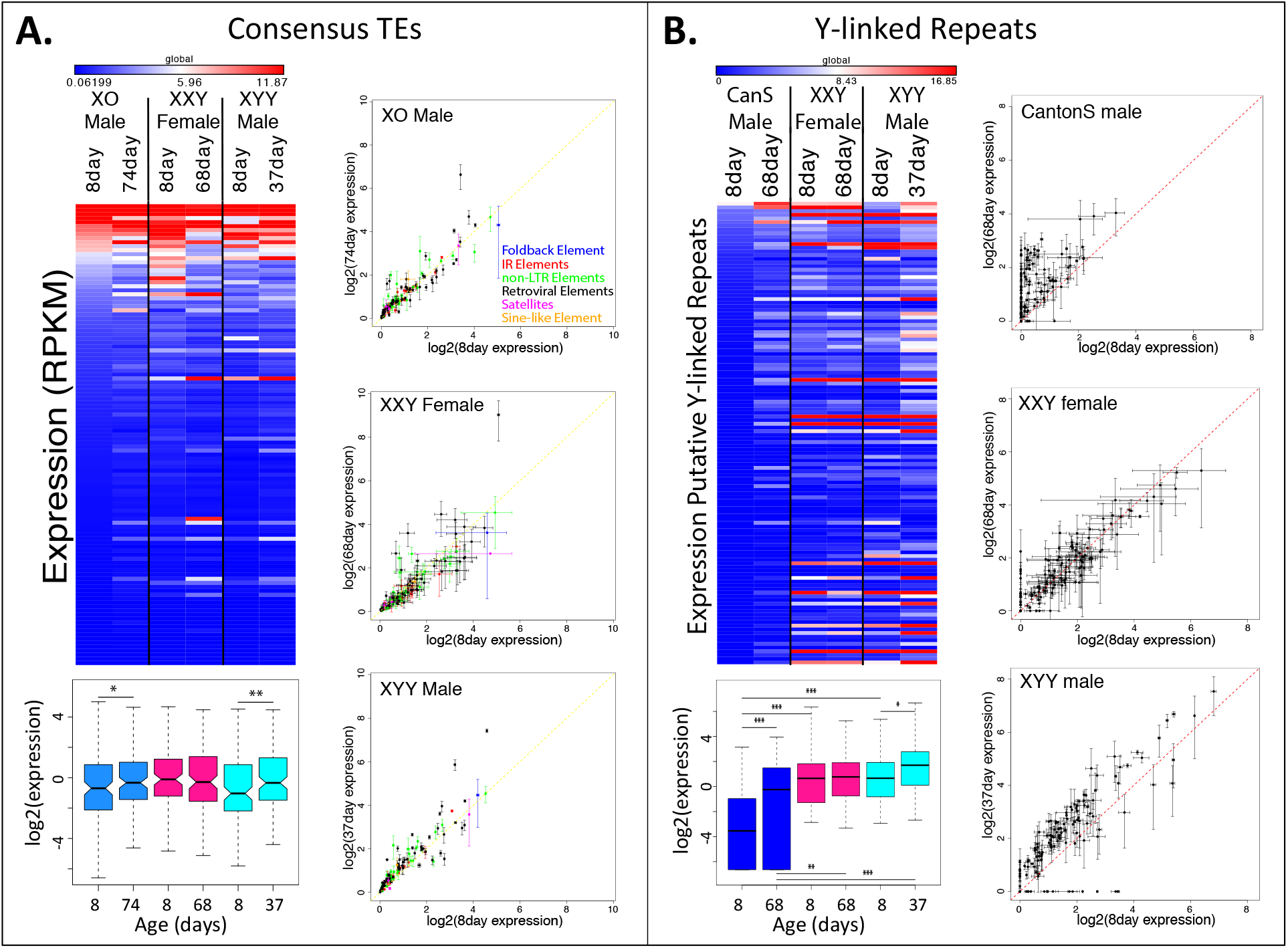
Expression of repetitive elements in XXY females and X0 and XYY males during aging. **A.** Expression of all repeats from FlyBase consensus library from the Release 6 of the *D. melanogaster* genome. The heatmap shows averaged expression across replicates, with significance values calculated using a Wilcoxon test (* p<0.1, ** p<0.05, *** p<0.01). The scatterplots show expression of all repeats from the FlyBase consensus library in young and old XO males, XXY females, and XYY males, with lines indicating the standard deviation of each expression value calculated from replicates, and color indicating the class of repetitive element. **B.** Expression for putatively Y-linked (male-specific) repeats in karyotypes with a Y chromosome, averaged across replicates, with significance calculated using the Wilcoxon test (* p<0.05, ** p<0.001, *** p<1e-5) Scatterplots like in **(A.)** for putatively Y-linked repeats.

Mis-expression of repetitive elements in XXY females and XYY males is especially pronounced for repeats found on the Y chromosome. Y-linked repeats show reduced silencing even in young XXY females and XYY males relative to wildtype males (**Figure 4B**) (Brown, et al. 2019), and become de-repressed even more during aging in XXY and XYY individuals (**Figure 4B**). Overall, wildtype males express 64 putatively Y-linked repeats during their lifespan, while XXY females express 86 and XYY males express 102. In XXY females, 13 repeats significantly increase in expression during aging (and 5 decrease), and 71 repeats significantly increase in expression in XYY males (and 8 decrease; **Figure S14**). Indeed, even at just 37 days old, XYY males already show higher, presumably aberrant expression of Y-linked repeats compared to 68-day-old wild-type males (**Figure 4B**).

## Discussion

Heterochromatin loss during aging has been observed in a large variety of species, ranging from yeast, to worms, flies and mammals (Haithcock, et al. 2005; Larson, et al. 2012; Zhang, et al. 2015). Here, we show that sex-specific heterochromatin loss contributes to sex-specific cellular aging in Drosophila. In particular, we find that increased heterochromatin loss in old male flies is associated with increased expression of repeats, and especially of repeats located on the Y chromosome. We further show that the absence or presence of a Y chromosome can directly influence lifespan in flies.

The molecular mechanisms underlying heterochromatin loss during aging are poorly understood, and our study does not allow us to address the causative cellular mechanisms underlying sex-specific heterochromatin loss and aging. One prominent heterochromatic structure that has been directly implicated in aging is the nucleolus (Ganley and Kobayashi 2014; Larson, et al. 2012; Lu, et al. 2018; Sinclair, et al. 1997). The nucleolus, the site of ribosome assembly, is formed at the tandemly repeated ribosomal DNA (rDNA) and is embedded in heterochromatin in most eukaryotes (Peng and Karpen 2007). In Drosophila, the rDNA cluster is located on the X and Y chromosome, each containing a few hundred rDNA transcriptional units in tandem repeats (Larson, et al. 2012; Lu, et al. 2018). Nucleolar size and activity have been mechanistically implicated in aging and longevity (Buchwalter and Hetzer 2017; Tiku, et al. 2017). In yeast, rDNA instability (i.e. reduction of rDNA copy number and associated accumulation of extrachromosomal rDNA circles) is a major cause of replicative aging (Ganley and Kobayashi 2014; Sinclair, et al. 1997). Destabilization and loss of rDNA has been shown during aging of male germline stem cells in Drosophila, which manifests cytologically as atypical morphology of the nucleolus (Lu, et al. 2018). In contrast, no such defects of nucleolar morphology were found in young and old female germline stem cells (Lu, et al. 2018). The rDNA locus is highly transcribed, and possible collusions between the replication and transcription machinery are thought to contribute to genomic instability of the rDNA (Helmrich, et al. 2013). In male Drosophila, transcription of rDNA is normally restricted to the Y chromosome (nucleolar dominance (Greil and Ahmad 2012)), and transcriptionally active Y-linked rDNA copies are preferentially lost in aging germline stem cells (Lu, et al. 2018). Flies with only a single rDNA cluster (i.e. X0 flies) live the longest, while flies with extra Y chromosomes (and thus more rDNA copies; see **Figure S16**) live shorter (**Figure 3**). This suggests a more complex picture on how the nucleolus may contribute to sex-specific aging, and future detailed functional experimentation is necessary to understand the molecular basis of the sex-specific differences in heterochromatin loss in *Drosophila*.

To conclude, our data demonstrate that the repeat-rich Y chromosome decreases life span in Drosophila. Loss of heterochromatin in repetitive regions during aging is more pronounced in male flies, and is accompanied by a de-repression of TEs. Y-linked repeats disproportionally lose their repressive marks and become reactivated, and analysis of flies with aberrant sex chromosome configurations demonstrates that the Y has a direct influence on organismal survival. Age-related heterochromatin loss on the repetitive, sex-limited Y or W-chromosome and repeat re-activation can contribute to lower survivorship of the heterogametic sex across taxa (Pipoly, et al. 2015), including humans. Y chromosomes of Drosophila and humans are known to harbor structural polymorphism in heterochromatic sequences and copy-number variation in repeats (Lyckegaard and Clark 1989; Repping, et al. 2003), and polymorphism on the *D. melanogaster* Y has been shown to affect lifespan (Griffin, et al. 2015), the formation of heterochromatin (Lemos, et al. 2010), and the regulation of TEs and 100s of genes genome-wide (Lemos, et al. 2008; Sackton, et al. 2011). More generally, individual humans and flies show extensive variation in their repeat content (Bosco, et al. 2007; Ewing and Kazazian 2010), and our results raise the question whether natural variation in repetitive sequences can contribute to genetic variation in longevity among individuals.

## Materials & Methods

### Drosophila strains

Fly strains were obtained from the Bloomington Stock Center and the Kyoto Stock Center. The following strains were used: Canton-S; Oregon-R; 2549 (C(1;Y),y^1^cv^1^v^1^B/0 & C(1)RM,y^1^v^1^/0); 4248 (C(1)RM, y^1^ pn^1^ v^1^ & C(1;Y)1, y^1^ B^1^/0; sv^spa-pol^) from the Bloomington Stock Center, and 100950 (0/ C(1)RM, y^1^w^str^/C(t;Y)1, y^1^ y^+^ ac^1^ sc^1^ w^1^) from the Kyoto Stock Center. The crossing scheme used to obtain XO and XYY males and XXY females is depicted in **Fig. 3A**. For chromatin and gene expression analyses, flies were grown in incubators at 25°C, 60% relative humidity, and 12h light for the indicated number of days following eclosion, and were then flash-frozen in liquid nitrogen and stored at −80°C. See **Fig. S17** for exact crossing scheme.

### Genome size estimation

We estimated genome size of the 5 karyotypes of interest using flow cytometry methods similar to those described in (Ellis, et al. 2014). Briefly, samples were prepared by using a 2mL Dounce to homogenize one head each from an internal control (*D. virilis* female, 1C=328 Mb) and one of the 5 karyotypes in Galbraith buffer (44mM magnesium chloride, 34mM sodium citrate, 0.1% (v/v) Triton X-100, 20mM MOPS, 1mg/mL RNAse I, pH 7.2). After homogenizing samples with 15-20 strokes, samples were filtered using a nylon mesh filter, and incubated on ice for 45 minutes in 25 ug/mL propidium iodide. Using a BD Biosciences LSR II flow cytometer, we measured 10,000 cells for each unknown and internal control sample. We ran samples at 10-60 events per second at 473 voltage using a PE laser at 488 nm. Fluorescence for each *D. melanogaster* karyotype was measured using the FACSDiva 6.2 software and recorded as the mode of the sample’s fluorescent peak interval. We calculated the genome size of the 5 karyotypes by multiplying the known genome size of *D. virilis* (328 Mb) by the ratio of the propidium iodide fluorescence in the unknown karyotype to the *D. virilis* control.

### Lifespan assays

Lifespan data was collected for all karyotypes in the same rearing conditions as described above. The lifespan assays were conducted as described (Linford, et al. 2013). Briefly, synchronized embryos were collected on agar plates, mobilized with a cotton swab, washed 3 times with PBS pH 7.4, and 10μl of embryos were pipetted to a fresh vial of standard fly medium. Adult flies were collected over 2 days, and were allowed to mate for 2 more days. Flies were then sexed, and 30 flies were counted into each vial. Vials were then flipped, without using CO_2_, every 2-3 days, and fly deaths were recorded. Flies that were observed escaping the vial were censored. To collect samples for the RNA-seq and ChIP-seq experiment, we censored the entire lifespan experiment once it reached 50% survivorship and flash-froze the remaining flies in liquid nitrogen. In total, 8,829 flies in 297 vials were counted for the lifespan assays reported here.

### Chromatin Immunoprecipitation and Sequencing

We performed ChIP-seq experiments using a standard protocol adapted from (Alekseyenko, et al. 2006). Briefly, approximately 2ml of adult flash-frozen flies were dissected on dry ice, and heads and thoraces were used to fix and isolate chromatin. Following chromatin isolation, we spiked in 60μl of chromatin prepared from female *Drosophila miranda* larvae (approximately 1μg of chromatin). We then performed immunoprecipitation using 4μl of the H3K9me2 (Abcam ab1220) antibody.

After reversing the cross-links and isolating DNA, we constructed sequencing libraries using the BIOO NextFlex sequencing kit. Sequencing was performed at the Vincent J. Coates Genomic Sequencing Laboratory at UC Berkeley, supported by NIH S10 Instrumentation Grants S10RR029668 and S10RR027303. We performed 50bp single-read sequencing for our input libraries, and 100bp paired-end sequencing for H3K9me2 libraries, due to their higher repeat content.

We collected replicate datasets for H3K9me2 enrichment in aged males and females to confirm differences seen between the sexes and between young and old samples (**Figure S3, S18**). Replicate H3K9me2 data for young flies are from (Brown, et al. 2019).

### RNA extraction and RNA-seq

We collected replicate RNA samples for aged individuals of all five karyotypes of interest; replicate RNA data for young flies are from (Brown, et al. 2019). Additionally, we collected 3 replicate samples for aged male Canton-S, aged XXY females, and aged XYY males, and 4 replicate samples for aged female Canton-S and aged XO males. After flash-freezing in liquid nitrogen, we dissected and pooled 5 heads from each sample, extracted RNA, and prepared stranded total RNA-seq libraries using Illumina’s TruSeq Stranded Total RNA Library Prep kit with Ribo-Zero ribosomal RNA reduction chemistry, which depletes the highly abundant ribosomal RNA transcripts (Illumina RS-122-2201). We performed single-read sequencing for all total RNA libraries at the Vincent J. Coates Genomic Sequencing Laboratory at UC Berkeley.

### Mapping of sequencing reads and data normalization

For all *D. melanogaster* alignments, we used Release 6 of the genome assembly and annotation (Hoskins, et al. 2015). For all ChIP-seq datasets, we used Bowtie2 (Langmead and Salzberg 2012) to map reads to the genome, using the parameters “-D 15 –R 2 –N 0 –L 22 –i S,1,0.50 --no-1-mm-upfront”, which allowed us to reduce cross-mapping to the *D. miranda* genome to approximately 2.5% of 50bp reads, and 1% of 100bp-paired end reads. We also mapped all ChIP-seq datasets to the *D. miranda* genome assembly (Ellison and Bachtrog 2013) to calculate the proportion of each library that originated from the spiked-in *D. miranda* chromatin versus the *D. melanogaster* sample.

To calculate the ChIP signal we first calculated the coverage across 5kb windows for both the ChIP and the input, and then normalized by the total library size, including reads that map to both *D. melanogaster* and the *D. miranda* spike. We then calculated the ratio of ChIP coverage to input coverage for each of the 5kb windows, and normalized by the ratio of *D. melanogaster* reads to *D. miranda* reads in the ChIP library, and then by the ratio of *D. melanogaster* reads to *D. miranda* reads in the input, to account for differences in the ratio of sample to spike present before immunoprecipitation. We describe the validation of this normalization method in (Brown, et al. 2019).

### Gene expression analysis

For each replicate of RNA-seq data, we first mapped RNA-seq reads to the ribosomal DNA scaffold in the Release 6 version of the *D. melanogaster* genome, and removed all reads that mapped to this scaffold, as differences in rRNA transcript abundance are likely to be technical artifacts from the total RNA library preparation, which aims to remove the bulk of rRNA transcripts. We then mapped the remaining reads to the Release 6 version of the *D. melanogaster* genome using Tophat2 (Kim, et al. 2013), using default parameters. We then used Cufflinks (Trapnell, et al. 2012) and Cuffdiff (Trapnell, et al. 2013) to merge replicates and calculate normalized FPKMs for all samples. GO analysis was performed using GOrilla, using ranked lists of differentially expressed genes (Eden, et al. 2009).

### Repeat libraries

We used two approaches to quantify expression of repeats. Our first approach was based on consensus sequences of known repetitive elements that were included in the Release 6 version of the *D. melanogaster* genome and are available on FlyBase. These included consensus sequences for 125 TEs and the 3 largest satellites (359, dodeca, and responder).

Our second approach aimed to specifically assess the repeat content of the Y chromosome. Since the Y chromosome is poorly assembled and repetitive elements on the Y are not annotated, we previously assembled repetitive elements *de novo* from male and female genomic DNA reads using RepARK and identified 101 male-specific repeats comprising 13.7kb of sequence, based on male-specific coverage analysis (Brown, et al. 2019; Koch, et al. 2014).

To assess expression of repetitive elements, we mapped RNA-seq reads to each of the repeat libraries (consensus TEs and putatively Y-linked repetitive elements) using Bowtie2 and the parameters “-D 15 –R 2 –N 0 –L 22 –i S,1,0.50 --no-1-mm-upfront”. We then calculated the mean coverage across each repetitive element using Bedtools, and normalized the coverage by the number of uniquely-mapping reads in the sequencing library. We made this calculation independently for each replicate for each time sample and karyotype, and then calculated both the average expression value as well as the standard deviation, to assess statistical significance and reproducibility (**Figure S19**).

To assess H3K9me2 signal in repetitive elements, we took a similar approach as we did for calculating ChIP enrichment profiles across the genome. First, we mapped both ChIP and input sequencing reads to each of the repeat libraries using Bowtie2 and the parameters “-D 15 –R 2 –N 0 –L 22 –i S,1,0.50 --no-1-mm-upfront”. We then calculated the mean coverage across each repetitive element using Bedtools, and normalized the coverage by the total library size, including reads that mapped to both the *D. melanogaster* and *D. miranda* genomes. We then calculated the ratio of ChIP coverage to input coverage for each repetitive element, and then normalized by the ratio of *D. melanogaster* reads to *D. miranda* reads in the ChIP library, and then by the ratio of *D. melanogaster* reads to *D. miranda* reads in the input, as described above and in (Brown, et al. 2019). This method accounts for differences in copy number of the repetitive elements by dividing the ChIP coverage by each repeat’s coverage in the input.

## Author contributions

DB and EB conceived the study and wrote the paper. EB collected and analyzed the data and AN analyzed data.

## Data availability

All RNA-seq and ChIP-seq reads are deposited in GeneBank (Accession number xx). The authors declare no competing financial interests. Correspondence and requests for materials should be addressed to.

## Supplementary information

**Figure S1.** Kaplan-Meier survivorship curves of line 2549 males and females ((C(1;Y),y^1^cv^1^v^1^B/0 & C(1)RM,y^1^v^1^/0) and Oregon-R wild-type males and females.

**Figure S2.** Same plot as Fig. 1C, but using a different normalization procedure (Bonhoure et al. 2014) (A), or using only uniquely mapping reads (B).

**Figure S3.** Pearson correlation coefficients for replicate H3K9me2 datasets for old males and females, and boxplots of normalized enrichment values for the replicates. Genome-wide plots were generated using replicate data as in Figure 1B. and 1D.

**Figure S4.** Chromosomal locations of 50kb windows that gain (red) or lose (blue) at least 1.5-fold H3K9me2 signal during aging for males and females. Pericentromeric regions are indicated by the red portion of the line beneath each chromosome.

**Figure S5.** Chromosomal locations of the top 10% of 50kb windows that gain (red) or lose (blue) H3K9me2 enrichment during aging for males and females. Pericentromeric regions are indicated by the red portion of the line beneath each chromosome.

**Figure S6.** Enrichment of H3K9me2 (in 5kb windows) for 1Mb upstream and downstream of the euchromatin/ pericentromere boundary, indicated by the dotted red line, on the 5 major chromosome arms. Subtraction plots show higher H3K9me2 signal in young (blue) or old (red) flies.

**Figure S7.** Expression values of all genes, normalized across replicates, of young and old males and females by chromosome location, as annotated in the Release 6 of the *D. melanogaster* genome. We consider expressed genes as those with FPKM>1, as determined by median intronic FPKM. Significance values are calculated using the Wilcoxon test.

**Figure S8.** Expression values of genes located in 50kb windows that show either a 1.5-fold loss or 1.5-fold gain of H3K9me2 during aging in males and females. Significance values are calculated using the Wilcoxon test.

**Figure S9.** Expression values of genes located in the top 10% of 50kb windows that either gain or lose H3K9me2 during aging in males and females. Significance values are calculated using the Wilcoxon test.

**Figure S10.** Overlap of the top 10% of differentially expressed genes during aging, normalized across replicates, for various combinations of the 5 sex chromosome karyotypes examined.

**Figure S11.** GO category enrichment of genes differentially expressed during aging in wild-type Canton-S males (**A**.) and Canton-S females (**B**.), and XO males (**C**.), XXY females (**D**.), and XYY males (**E**.). Genes were ranked by their fold change in expression, averaged across replicates, regardless of direction, and submitted to GOrilla (Eden, et al. 2009) for GO category enrichment analysis.

**Figure S12.** Number of repeats that show a significant increase (red) or decrease (blue) in expression during aging as a fraction of all repeats from the FlyBase consensus repeat library, with significance estimated using standard errors from replicate datasets. Significance is calculated using Fisher’s exact test, with red stars indicating significance for repeats that increase in expression, and blue stars indicating significance for repeats that decrease in expression during aging. We also show the estimates of the total fraction of RNA-seq reads that map to the FlyBase consensus repeat library, with error bars calculated from replicate datasets, for young and old samples from each of the 5 karyotypes.

**Figure S13.** Male vs. female genomic coverage of *de novo* assembled repeats, with putatively Y-linked repeats indicated in blue and purple as those with male-specific or highly male-biased genomic coverage patterns (Brown, et al. 2019).

**Figure S14.** Number of putative Y-linked repeats that show a significant increase (red) or decrease (blue) in expression during aging as a fraction of all repeats from a male-specific repeat library (see **Figure S12**), with significance estimated using standard errors from replicate datasets. Significance is calculated using Fisher’s exact test, with red stars indicating significance for repeats that increase in expression, and blue stars indicating significance for repeats that decrease in expression during aging. We also show the estimates of the total fraction of RNA-seq reads that map to the Y-specific consensus repeat library, with error bars calculated from replicate datasets, for young and old samples from each of the 5 karyotypes.

**Figure S15.** Expression values of all genes, normalized across replicates, of young and old XO males, XXY females, and XYY males by chromosome location, as annotated in the Release 6 of the *D. melanogaster* genome. Significance values are calculated using the Wilcoxon test.

**Figure S16.** Genomic coverage across the rDNA locus for flies with different sex chromosome karyotypes. Coverage is normalized to autosomes, to roughly infer rDNA copy number. The numbers show median coverage across the rDNA locus.

**Figure S17.** Crossing scheme to generate flies with aberrant sex chromosomes.

**Figure S18.** Estimates of H3K9me2 signal for the FlyBase consensus library for all replicates for all karyotypes.

**Figure S19.** Estimates of expression values for the FlyBase consensus library for all replicates for all karyotypes.

**Table S1.** Average estimation, across all replicates, of the fraction of all RNA-seq reads that are derived from the FlyBase consensus repeat library, as well as the fold change in repetitive content during aging.

**Table S2.** Average estimation, across all replicates, of the fraction of all RNA-seq reads that are derived from the putative Y-linked consensus repeat library, as well as the fold change in repetitive content during aging.

